# The *C. elegans* homolog of *TMEM132D*, a human panic-disorder and anxiety risk gene, modulates neuronal morphogenesis through the WAVE-regulatory complex

**DOI:** 10.1101/2020.11.16.385906

**Authors:** Xin Wang, Wei Jiang, Shuo Luo, Xiaoyu Yang, Changnan Wang, Bingying Wang, Yongjun Dang, Yin Shen, Dengke K. Ma

## Abstract

*TMEM132D* is a human gene identified with multiple risk alleles for panic disorders, anxiety and major depressive disorders. Belonging to a conserved family of transmembrane proteins, TMEM132D and its homologs are still of unknown molecular functions. By generating loss-of-function mutants of the sole TMEM132 ortholog in *C. elegans*, we identify abnormal morphologic phenotypes in the dopaminergic PDE neurons. Using a yeast two-hybrid screen, we find that NAP1 directly interacts with the cytoplasmic domain of human TMEM132D, and mutations in *C. elegans tmem-132* that disrupt the interaction with NAP1 cause similar morphologic defects in the PDE neurons. NAP1 is a component of the WAVE regulatory complex (WRC) that controls F-actin cytoskeletal dynamics. Decreasing activity of WRC rescues the PDE defects in *tmem-132* mutants, whereas gain-of-function of *TMEM132D* in mammalian cells inhibits WRC, leading to decreased abundance of selective WRC components, impaired actin nucleation and cell motility. We propose that metazoan TMEM132 family proteins play evolutionarily conserved roles in regulating NAP1 protein homologs to restrict inappropriate WRC activity, cytoskeletal and morphologic changes in the cell.

## INTRODUCTION

Despite decades of genetic and molecular analyses, the genome of the common model organism *C. elegans* still comprises many functionally uncharacterized genes (*C. elegans* Sequencing Consortium, 1998; Hillier et al., 2005; Kim et al., 2018; Pandey et al., 2014). One such example is the *C. elegans* gene *Y71H2AM.10*, an ortholog of the evolutionarily conserved *TMEM132* gene family (Kim et al., 2018; Sanchez-Pulido and Ponting, 2018). This family encodes single-pass transmembrane proteins present in metazoans but remains functionally uncharacterized in any organisms. The human genome encodes 5 paralogs (*TMEM132A-E*), genetic variants of which have been identified as risk alleles of many human diseases, including those associated with panic disorder and anxiety severity in *TMEM132D* (Erhardt et al., 2011, 2012; Hodgson et al., 2016; Howe et al., 2016; Inoue et al., 2015; Quast et al., 2012; Shimada-Sugimoto et al., 2016). Risk variants of *TMEM132D* have been shown to correlate with altered mRNA levels of *TMEM132D* in anxiety-related brain regions and psychiatric syndromes (Erhardt et al., 2011; Howe et al., 2016). In addition, the *TMEM132D* locus in cattle appears to have undergone an evolutionary selective sweep during domestication, along with reduced fearfulness in cattle’s behaviors (Qanbari et al., 2014). The expression of *TMEM132* family genes is also highly enriched in the nervous system of diverse animals, including *C. elegans* and humans (Cao et al., 2017; Fagerberg et al., 2014). However, it remains unknown how TMEM132 family proteins regulate neuronal structure and function and how their abnormal function and regulation may contribute to various neurological and psychiatric diseases.

Neuronal morphological changes are driven primarily by actin cytoskeletal dynamics under the control of the WAVE-regulatory complex (WRC). WRC promotes actin nucleation to form filamentous actin (F-actin) by stimulating activity of the Arp2/3 complex in response to biochemical signals originating from a variety of neuronal membrane receptors (Chen et al., 2014; Chia et al., 2014; Eden et al., 2002). WRC is a multi-subunit complex comprising SRA1, HSPC300, ABI1/2, WAVE1/2/3 and NAP1 (also known as NCKAP1) proteins. *NAP1* was initially identified as a gene with strongly decreased expression in the brain of patients with sporadic Alzheimer’s disease (Suzuki et al., 2000). Deleterious *NAP1* variants were also identified in human patients with autism and intellectual disability (Anazi et al., 2017; Freed and Pevsner, 2016). Among other components in the WRC, the SRA and ABI proteins form an evolutionarily conserved binding interface for diverse WRC ligands that commonly contain the WRC-interacting receptor sequence (WIRS) motif (Chen et al., 2014; Chia et al., 2014). Whether NAP1 directly interacts with any neuronal membrane receptors to affect WRC signaling and actin cytoskeletal changes has not been reported. It has also been unclear how abundance of WRC components is regulated in cell compartments where actin nucleation needs to be limited in morphologically complex cells, including neurons.

To elucidate biological functions and mechanisms of action of TMEM132 family proteins, we generated and characterized *C. elegans* loss-of-function (LOF) mutations in *Y71H2AM.10*, the sole ortholog of the *TMEM132* gene family. *tmem-132* mutants exhibit striking morphological defects in the dopaminergic PDE neurons. We further identified human NAP1 as a TMEM132D interactor and show that the *C. elegans* homologs also interact with each other. Genetic interactions between *tmem-132* and WRC-encoding genes in regulating PDE morphology, the LOF phenotype of *tmem-132* mutants and gain-of-function (GOF) phenotype of *TMEM132D* in mammalian cells collectively suggest that TMEM132 family proteins regulate NAP1 levels in WRC to finely modulate actin nucleation, cellular cytoskeletal and morphological changes.

## RESULTS

### The *C. elegans* TMEM-132 localizes to neurons and regulates the dopaminergic PDE neuron morphology

Protein homology and motif analysis identified the Pfam16070 domain characteristic of both human TMEM132D and *C. elegans* TMEM-132, classifying both to the evolutionarily conserved TMEM132 protein superfamily (Figure 1A; Figure S1). Interestingly, a translational reporter that fuses *tmem-132* with GFP revealed enriched expression of *C. elegans tmem-132* in neurons (Figure 1B-D), implicating a specific role of TMEM-132 in the nervous system. Lack of mammalian loss-of-function models and potential genetic redundancy among the TMEM132A-E family members precluded us from analyzing the physiological function of TMEM132D *in vivo*. Thus, we sought to address this issue in *C. elegans*, which encodes *tmem-132* as the sole ortholog of the gene family and has served as excellent model to study neuronal cell biology (Inberg et al., 2019; Richardson and Shen, 2019; Tang and Jin, 2018). We used CRISPR-Cas9 techniques to generate a series of *C. elegans* mutants, including multiple independently-derived deletions, an early stop-codon mutant and a genetic P784T knock-in mutant, in which the highly conserved proline residue became threonine, corresponding to the human disease risk allele for anxiety and panic disorders (Figures 1E and 1F). We generated such multiple independent mutations to seek convergent phenotype and outcrossed all mutants to eliminate potential interference of phenotype by background mutations.

**Figure 1.**
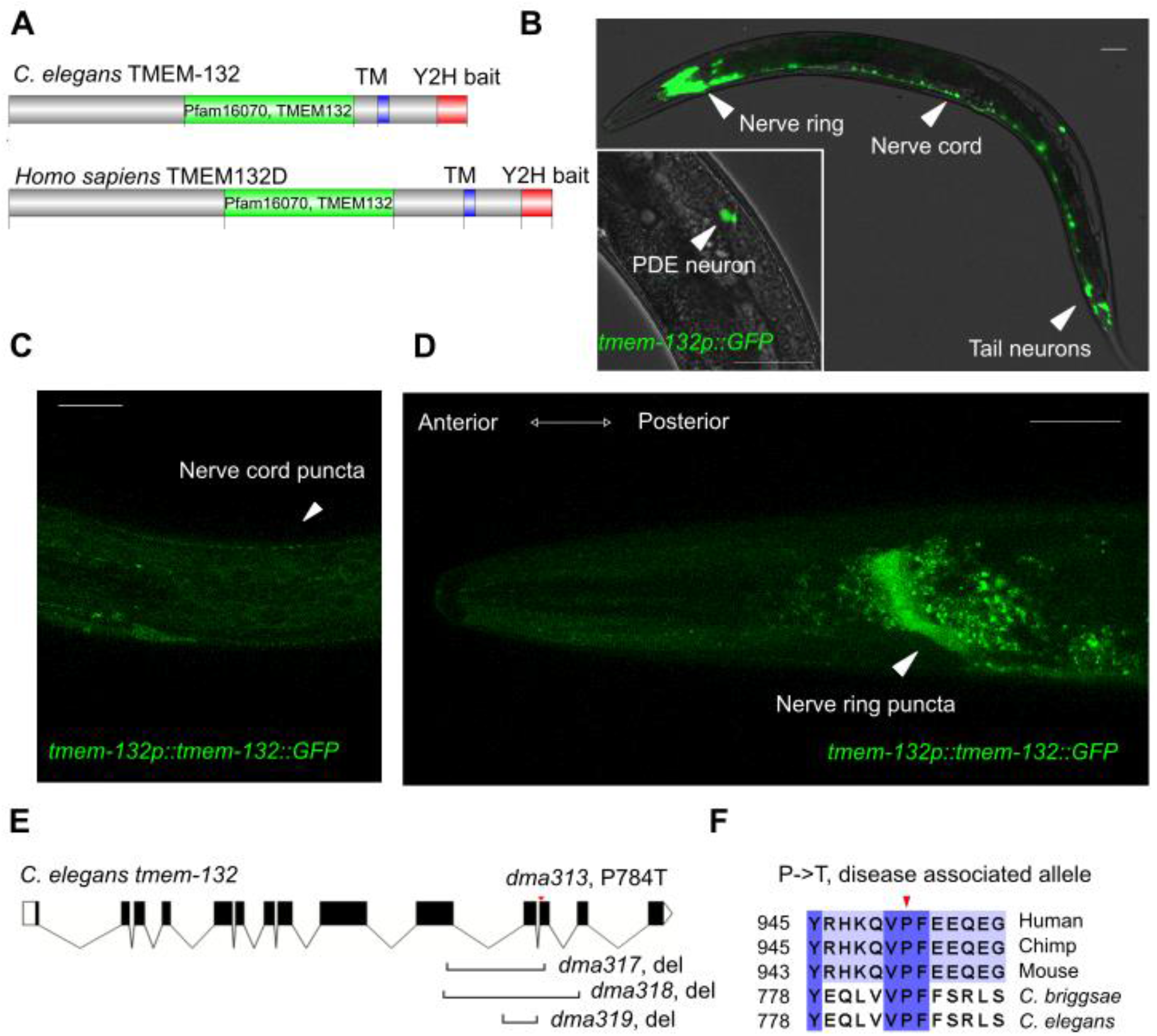
*C. elegans* TMEM-132 specifically localizes to the nervous system. **(A)** Schematic of human TMEM132D and *C. elegans* TMEM-132 protein domains. The Pfam16070 conserved domain (yellow) characterizes both proteins, with additional transmembrane domains (TM, blue) and C-terminal regions (red) used for yeast-two-hybrid assays. **(B)** Exemplar confocal fluorescence image showing expression of *tmem-132* promoter-driven GFP in the nervous system. Inset, exemplar confocal fluorescence image showing expression of *tmem-132* promoter-driven GFP specifically in the PDE neuron. **(C)** Exemplar confocal fluorescence image showing the puncta pattern of *tmem-132* promoter-driven TMEM-132∷GFP expression in the ventral nerve cord. **(D)** Exemplar confocal fluorescence image showing the puncta pattern of *tmem-132* promoter-driven TMEM-132∷GFP expression in the nerve ring. Scale bar: 50 μm. **(E)** Schematic of *tmem-132* gene structure showing positions of alleles generated by CRISPR-mediated editing including deletion alleles *dma317*, *dma318*, *dma319* and point mutation *dma313* that converts proline 784 to threonine. **(F)** Multiple sequence alignment (generated by ClustalOmega and visualized by Jalview) of TMEM132 protein family from indicated metazoan species showing high levels of amino acid sequence conservation around the P784 position. P784T is one of the non-synonymous variants identified as risk alleles for panic disorder and anxiety.

Given the exclusive localization of TMEM-132 in neurons, we subjected *tmem-132* mutants to a variety of neuronal phenotypic analyses. We did not observe gross behavioral defects under normal conditions. We next crossed the mutants to various established GFP reporters to examine neurons of stereotyped morphology, including ciliated sensory AWC neurons, hermaphrodyte-specific HSN neurons, mechanosensory PVD, ADE and PDE neurons (Figure S2). Among the neurons examined, the dopaminergic neuron PDE exhibited the most severe defect, thus in this study we focused on PDE, which is marked by the *osm-6*p∷GFP reporter in addition to other ciliated and dopaminergic neurons. Although dense GFP signals prevented us from analyzing the anterior group of ciliated and dopaminergic neurons, close confocal microscopic analysis of the posterior, anatomically isolated PDE neurons revealed striking abnormal morphologies in a large fraction of *tmem-132* mutants (Figures 2A-D). We categorized the mutant phenotype into several classes, including those with irregular soma outlines, ectopic dendrites, ectopic axon branches, and axon misguidance as similarly described previously (Shakir et al., 2008; Sulston et al., 1975). Although the phenotypic defects of PDE are diverse, all mutants show similar profiles in distribution of different categories of phenotypic defects (Figures 2E and 2F). *tmem-132* LOF mutants also exhibited morphological defects in the ADE and PVD but not morphologically less complex AWC neurons (Figure S2). Neuronal morphogenesis critically depends on neuronal interactions with glia and epithelia in *C. elegans* (Inberg et al., 2019; Lamkin and Heiman, 2017; Singhvi and Shaham, 2019). Importantly, transgenic expression of *tmem-132* driven by the *osm-6* promoter rescued the morphologic defect of *tmem-132* mutants, indicating a causal and cell-autonomous role of TMEM-132 for ensuring normal PDE neuronal morphology (Figure 2F).

**Figure 2.**
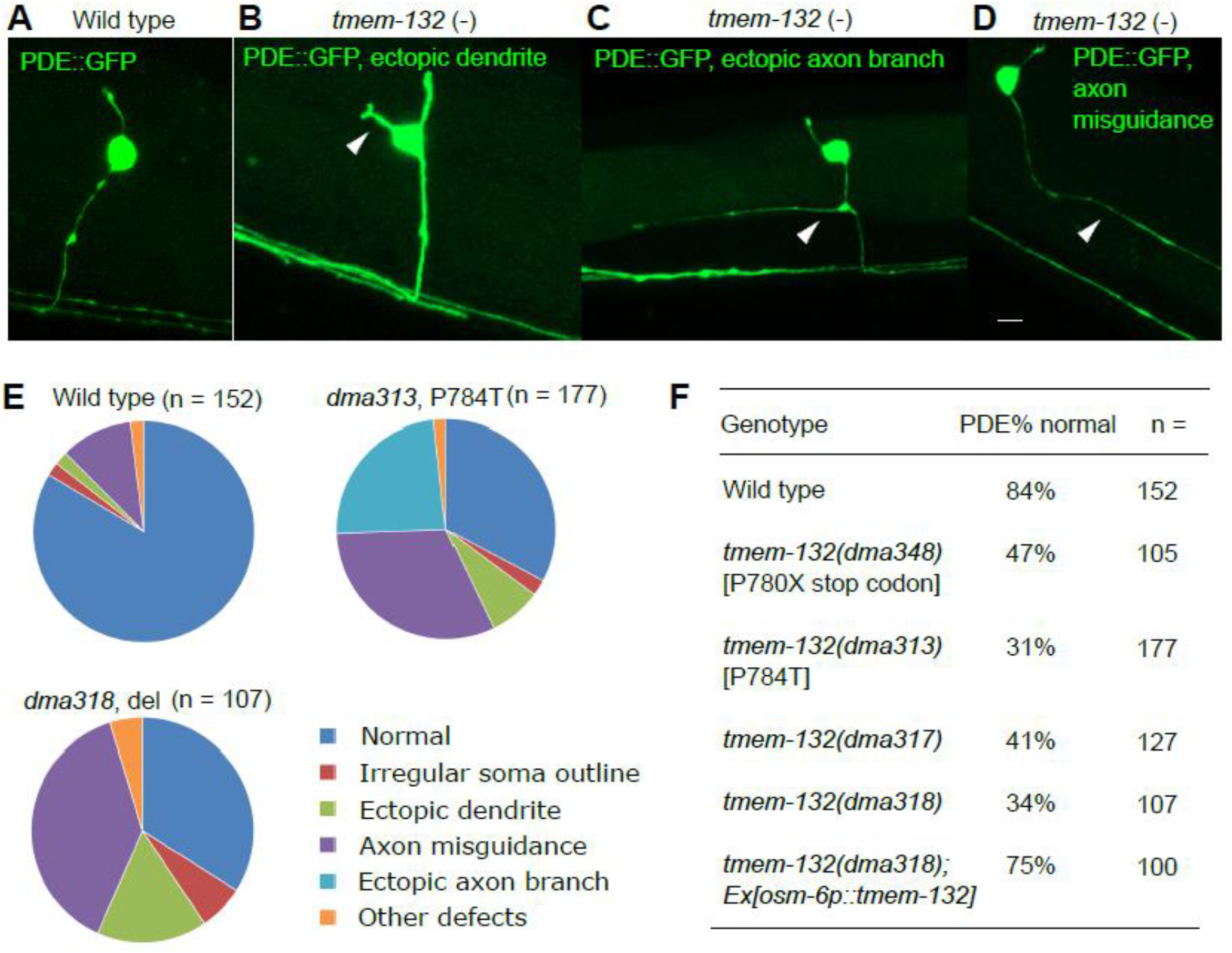
*C. elegans* TMEM-132 is crucial for correct morphology of the PDE neuron. **(A to D)** Confocal fluorescence images showing indicated major categories of abnormal morphology of PDE neurons in *tmem-132* deficient *C. elegans*. **(E)** Quantification of percentage of animals with abnormal PDE neuronal morphology in each indicated category shown in pie charts. **(F)** Table summarizing overall percentage of animals with normal PDE neuronal morphology in wild type and *tmem-132* mutants, including those carrying three independent deletion alleles, the P784T knock-in allele and the Q781X nonsense allele that truncates the C-terminus of TMEM-132. All strains carry the PDE reporter *osm-6*p∷GFP and were outcrossed to minimize potential effects of background mutations. Scale bars: 5 μm.

### Human TMEM132D interacts with the WRC component NAP1

To begin to understand molecular functions of this conserved protein superfamily, we used yeast-two-hybrid (Y2H) screens to identify protein interactors of human TMEM132D. The predicted intracellular C-terminus of TMEM132D contains cytoplasmic motifs likely related to actin cytoskeletal dynamics (Chen et al., 2014; Sanchez-Pulido and Ponting, 2018). We thus constructed a yeast bait vector expressing its C-terminal domain and used the bait to screen for interactors from a human-cDNA prey library. We also constructed a bait vector that contains the homologous C-terminus of *C. elegans tmem-132* and focused on identified screen hits whose homologs can interact with human and *C. elegans* baits, respectively. From 117 independent cDNA clones isolated and identified by Sanger sequencing (Table S1), we found that the protein NAP1 encoded by *NCKAP1* showed robust interaction with TMEM132D in Y2H assays (Figure 3A). NAP1 is an integral component of the WAVE regulatory complex that regulates actin nucleation and cytoskeletal changes in the cell through the ARP2/3 complex (Chen et al., 2014, 2010; Eden et al., 2002; Welch and Mullins, 2002). We found that GEX-3, the *C. elegans* ortholog of NAP1, also interacts with the C-terminus of *C. elegans* TMEM-132 (Figure 3A).

**Figure 3.**
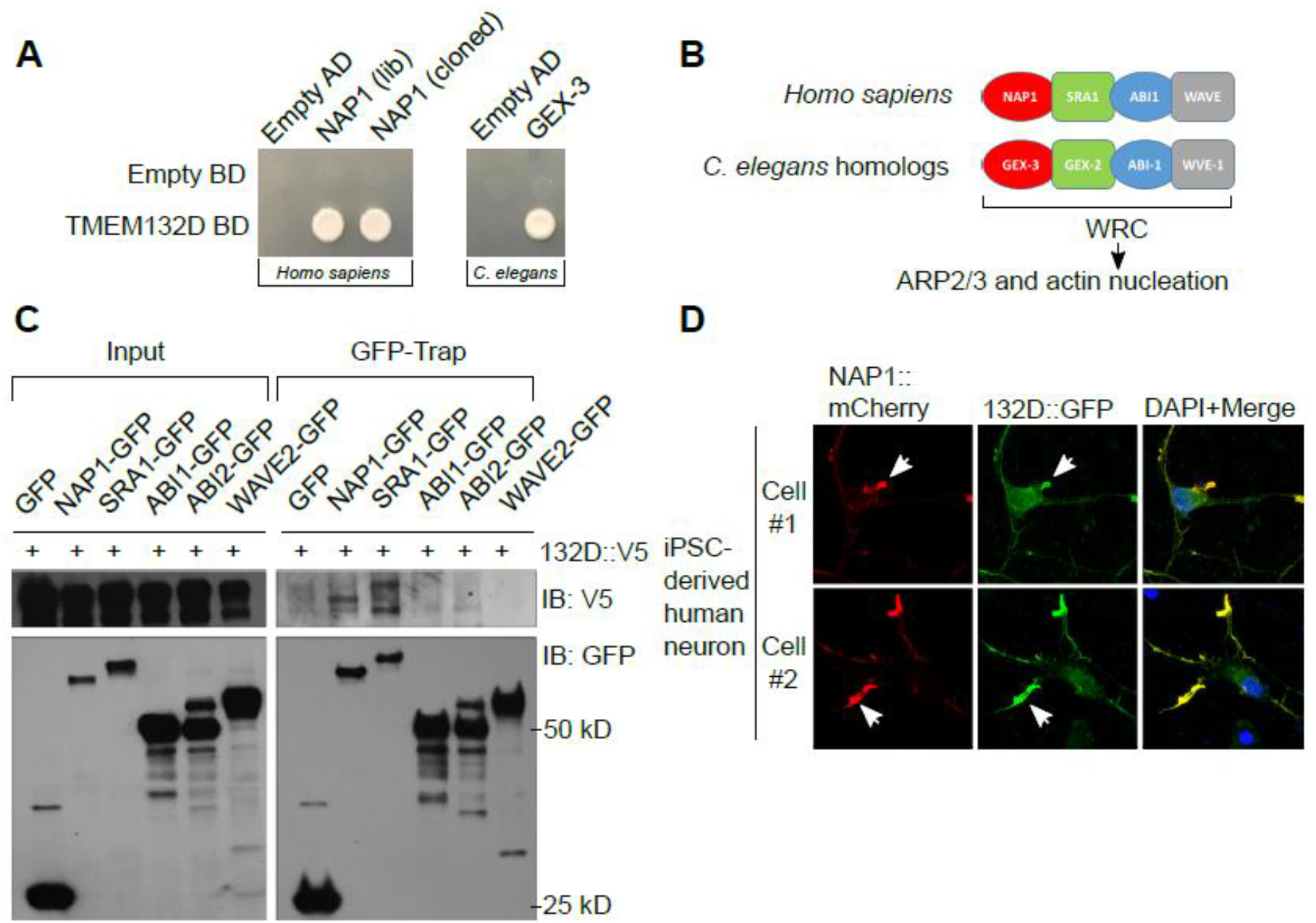
Conserved interactions between TMEM132 and WRC in human and *C. elegans*. **(A)** Yeast colony growth indicating interaction between protein C-termini from human TMEM132D and NAP1, as well as *C. elegans* TMEM-132 and GEX-3. Both library-derived and re-cloned NAP1 cDNAs showed specific interaction with TMEM132D. BD and AD refer to bait/prey vectors as controls. **(B)** Schematic of human and *C. elegans* homologs encoding various major components of WRC that controls actin nucleation via the ARP2/3 complex. **(C)** Exemplar Western blot showing biochemical interaction between V5-tagged TMEM132D and GFP-tagged NAP1 in CoIP assay. Mammalian expression vectors encoding TMEM132D-V5 and GFP-tagged WRC components were co-transfected to HEK293 cells followed by GFP-Trap CoIP and blotting with antibodies against V5. Only NAP1 and SRA1 showed robust interaction with TMEM132D. **(D)** Exemplar confocal immunofluorescence images showing co-localization of GFP-tagged TMEM132D and mCherry-tagged NAP1 in iPSC-derived human neurons transfected with both fluorescence reporters. Scale bar, 10 μm.

In addition to NAP1, the WRC also contains three other major proteins SRA1, ABI1/2 and WAVE1/2/3, with corresponding orthologs *gex-2*, *abi-1* and *wve-1* in *C. elegans* (Figure 3B) (Chia et al., 2014; Shakir et al., 2008; Soto et al., 2002). We verified the biochemical interaction between full-length NAP1 and TMEM132D in mammalian cells by co-immunoprecipitation (CoIP) assays. GFP-tagged NAP1, when expressed in heterologous HEK293 cells with V5 epitope-tagged TMEM132D, was able to pull down TMEM132D in CoIP (Figure 3C). Another component of WRC, SRA1, is structurally similar to NAP1 and together with NAP1 forms a heterodimeric sub-complex in WRC (Chen et al., 2010). GFP-tagged SRA1 also pulled down TMEM132D, although we did not observe apparent association of TMEM132D with other components of WRC. When co-expressed in fully differentiated human neurons derived from induced pluripotent stem cells (iPSC), GFP-tagged TMEM132D markedly co-localized with mCherry-tagged NAP1 (Figure 3D). Collectively, these genetic, biochemical and cellular imaging results identify NAP1 as a protein interactor of TMEM132D and indicate that such interaction is evolutionarily conserved also for *C. elegans* counterparts.

### The C-terminal domain and the conserved P784 are required for the interactions between TMEM132 and WRC components

We used Y2H and CoIP assays to further define the C-terminal domain of human TMEM132D or *C. elegans* TMEM-132 that is crucial for interacting with C-termini of NAP1 homologs. To examine interaction between GEX-3 (*C. elegans* NAP1 homolog) and TMEM-132, we generated transgenic strains in which HA epitope-tagged GEX-3 and mCherry-tagged TMEM-132 Ct are co-induced by heat shock promoters (Figure 4A). Using the mCherry nanobody-Trap CoIP assay, we found that mCherry-tagged TMEM-132 Ct specifically pulled down HA epitope-tagged GEX-3, compared with heat shock-induced mCherry only as control (Figure 4A). We also confirmed interaction between GEX-3 Ct and TMEM-132 Ct in Y2H assays, in which the most C-terminal 60 a.a. of GEX-3 was sufficient to mediate the interaction (Figure 4B). In the C-terminus homologous to the WIRS-containing TMEM132D, mutation of a WIRS-like motif attenuated TMEM-132 interaction with GEX-3 (Figure 4B). Furthermore, we generated mutations to convert the conserved proline 784 to alanine or threonine (to model psychiatric disorder-associated risk allele in humans, see Figure 1F) in TMEM-132 and found that both mutations abolished the interaction with GEX-3 (Figure 4C).

**Figure 4.**
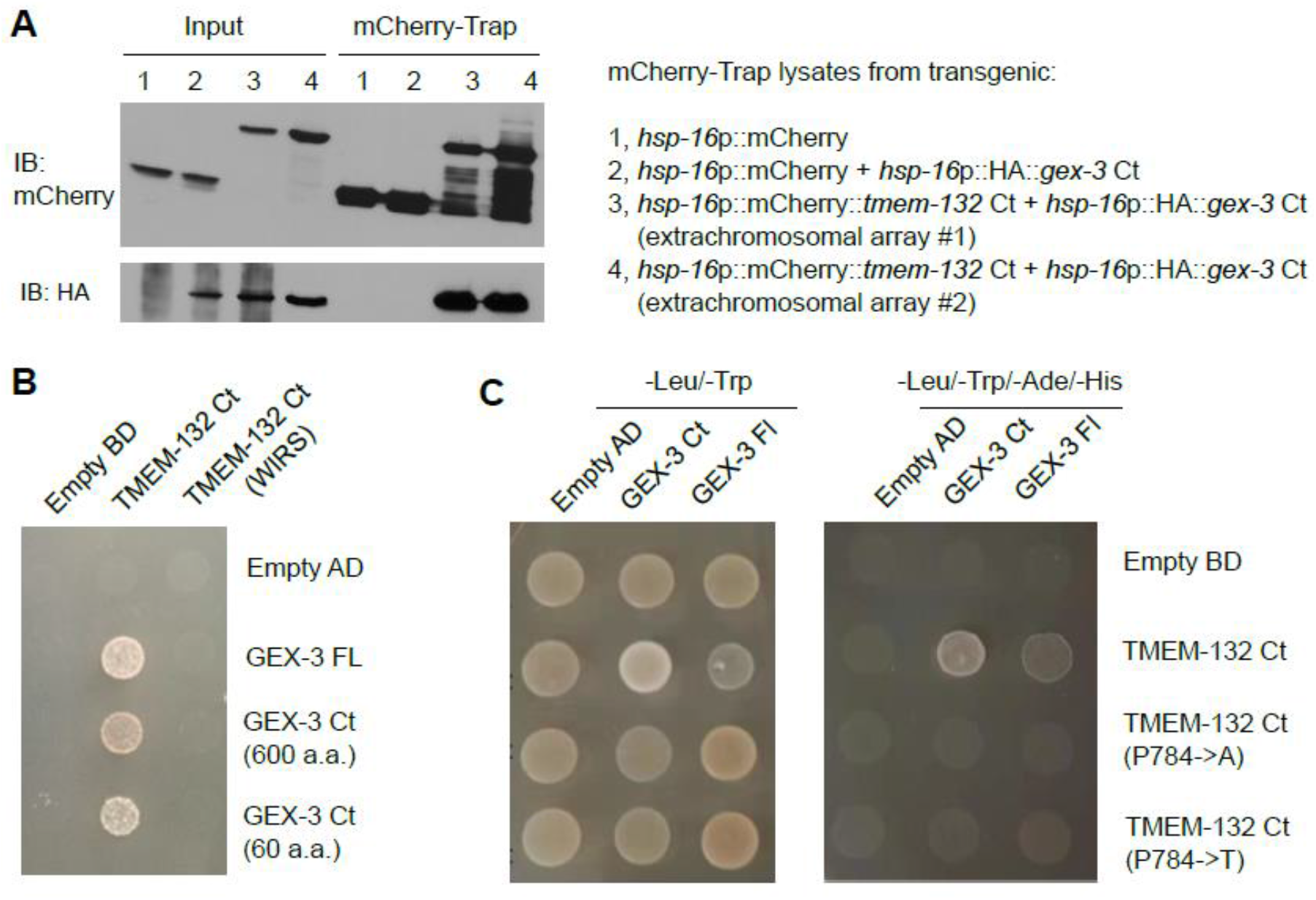
*C. elegans* TMEM-132 interaction with GEX-3 requires C-terminal domains involving key amino acid residues. **(A)** Exemplar Western blot showing CoIP of *C. elegans* TMEM-132 (mCherry-tagged) and GEX-3 (HA-tagged) C terminal domains. Transgenic expression was driven by *hsp-16* promoters that are heat shock inducible by placing transgenic animals at 32 °C for 2 hrs followed by recovery at 20 °C for 4 hrs. **(B)** Yeast growth colonies showing interaction of *C. elegans* TMEM-132 Ct and a mutant with a putative WIRS-like motif converted to alanine residues, with GEX-3 full length, mutants with C-terminal 600 a.a. and 60 a.a. fragments. **(C)** Yeast growth colonies showing interaction of *C. elegans* GEX-3 full length, mutants with C-terminal 600 a.a. fragments with TMEM-132 Ct and mutants with substitutions of proline 784 to alanine or threonine, respectively. Double dropout -Leu/-Trp yielded colonies without apparent differences, indicating that these mutations do not affect protein levels.

Previous studies revealed that the WIRS of diverse transmembrane proteins in mammals mediates binding to an interaction surface of WRC (Chen et al., 2014). As TMEM132D contains such a motif at its C-terminus, we performed mutation analysis in Co-IP assays and found that deletion of the entire cytoplasmic portion, deletion of the 120 a.a. C-terminus or mutation of the WIRS-like motif in TMEM132D markedly attenuated its interaction with Nap1 (Figure 5A). Deletion of the C-terminal 60 a.a. of TMEM132D did not appear to affect the interaction, indicating that additional sites beyond the 60 a.a. may interact with WIRS and contribute to interaction with Nap1. We made similar observations for Sra1 (Figure 5B), consistent with previous structural findings that Nap1 and Sra1 form an integral heterodimeric sub-complex of WRC (Chen et al., 2010). Systematic deletion mutation analysis using Co-IP and Y2H assays underscored the importance of C termini of TMEM132 family proteins from both *C. elegans* and humans in interacting with Nap homologs (Figure 5C). Since canonical WIRS binds to a composite surface formed by Sra and Abi but not Nap (Chen et al., 2014), our results suggest that TMEM132 differs from certain canonical WRC ligands, such as PCDH10, in specific interaction with WRC components, consistent with the idea that TMEM132 acts to sequester selective components of WRC rather than to recruit or activate WRC.

**Figure 5.**
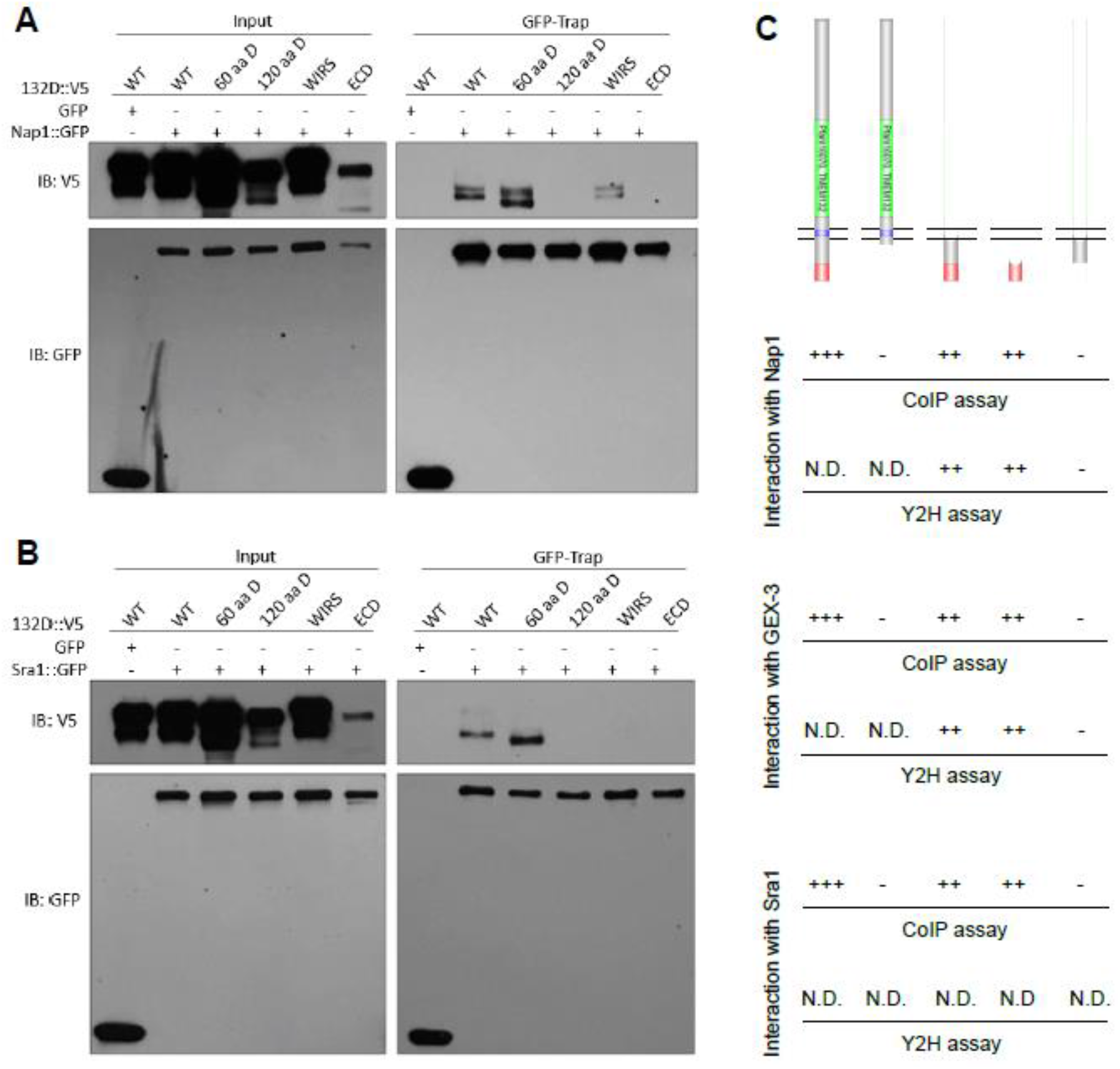
Human TMEM132D interacts with Nap/Sra via C-terminal domains. **(A)** Exemplar Western blot showing CoIP of various human TMEM132D (V5-tagged) mutants and NAP1 (GFP-tagged). Expression was driven by constitutive CMV promoters in HEK293 cells and lysates were used for GFP-trap and blot with antibodies against GFP and V5. **(B)** Exemplar Western blot showing CoIP of various human TMEM132D (V5-tagged) mutants and SRA1 (GFP-tagged). 60 a.a. D, TMEM132D with deletion of the last 60 a.a. sequence. 120 a.a. D, TMEM132D with deletion of the last 120 a.a. sequence. WIRS, TMEM132D with the WIRS-like motif (KFTTFTAV) mutated to alanine residues. ECD, TMEM132D with only extracellular domain and transmembrane domain. Results represent three independently repeated experiments. **(C)** Schematic showing human TMEM132D or *C. elegans* TMEM-132 with various domain genetic deletions and a summary of their interaction with Nap homologs (human NAP1 and *C. elegans* GEX-3) from both CoIP and Y2H assays.

### WRC acts downstream of TMEM-132 to regulate morphology of the PDE neurons

Since *C. elegans* TMEM-132 binds to GEX-3 as human TMEM132D binds to NAP1, we next addressed whether TMEM-132 regulates neuronal morphology via WRC in *C. elegans*. We generated an integrated transgenic reporter with neuronal specific expression of the WRC component ABI-1 fused to GFP. We found that ABI-1∷GFP in the nerve ring, along with the ganglia of the head and tail in *C. elegans*, is weakly fluorescent in wild type animals but strongly up-regulated in *tmem-132* mutants (Figures 6A-C). Close microscopic analysis of ABI-1∷GFP specifically in PDE neurons revealed that ABI-1∷GFP forms puncta, numbers of which decrease from the larval to adult stages in wild type animals (Figure 6D). By contrast, numbers of ABI-1∷GFP puncta in *tmem-132* mutants remain high in both larval and adult stages. Increased abundance of ABI-1∷GFP in *tmem-132* mutants were confirmed by both whole-animal Western blot and quantitative phenotypic penetrance analysis (Figures 6A-E). To assess whether abnormally high ABI-1∷GFP abundance in *tmem-132* mutants is responsible for morphologic defects of PDE neurons, we fed *tmem-132* mutants with bacteria expressing double-stranded RNAi against *abi-1* and found that morphologic defects of PDE neurons were largely normalized (Figures 6F and S3). RNAi by feeding produces weaker loss-of-function effects in neurons than by genetic deletion of WRC component-encoding genes, which by itself can cause strong PDE morphologic defects (Shakir et al., 2008). RNAi against genes encoding other components of WRC, including *brk-1* and *wve-1*, also normalized defects of PDE neurons (Figure 6G), supporting that abnormally high WRC activity in *tmem-132* mutants caused PDE defects. Together, these results indicate that TMEM-132 ensures normal PDE morphology by regulating the neuronal abundance of ABI-1 and restricting WRC activity in the PDE neuron. Interestingly, the CRISPR-mediated P784T substitution in the endogenous TMEM-132 locus caused abnormal neuronal morphology of the PDE neurons in *C. elegans* (Figure 2F), supporting the notion that functional roles of TMEM-132 in regulating F-actin and cell morphological changes are mediated by its C-terminal interaction with GEX-3 and thereby interference of WRC and actin nucleation.

**Figure 6.**
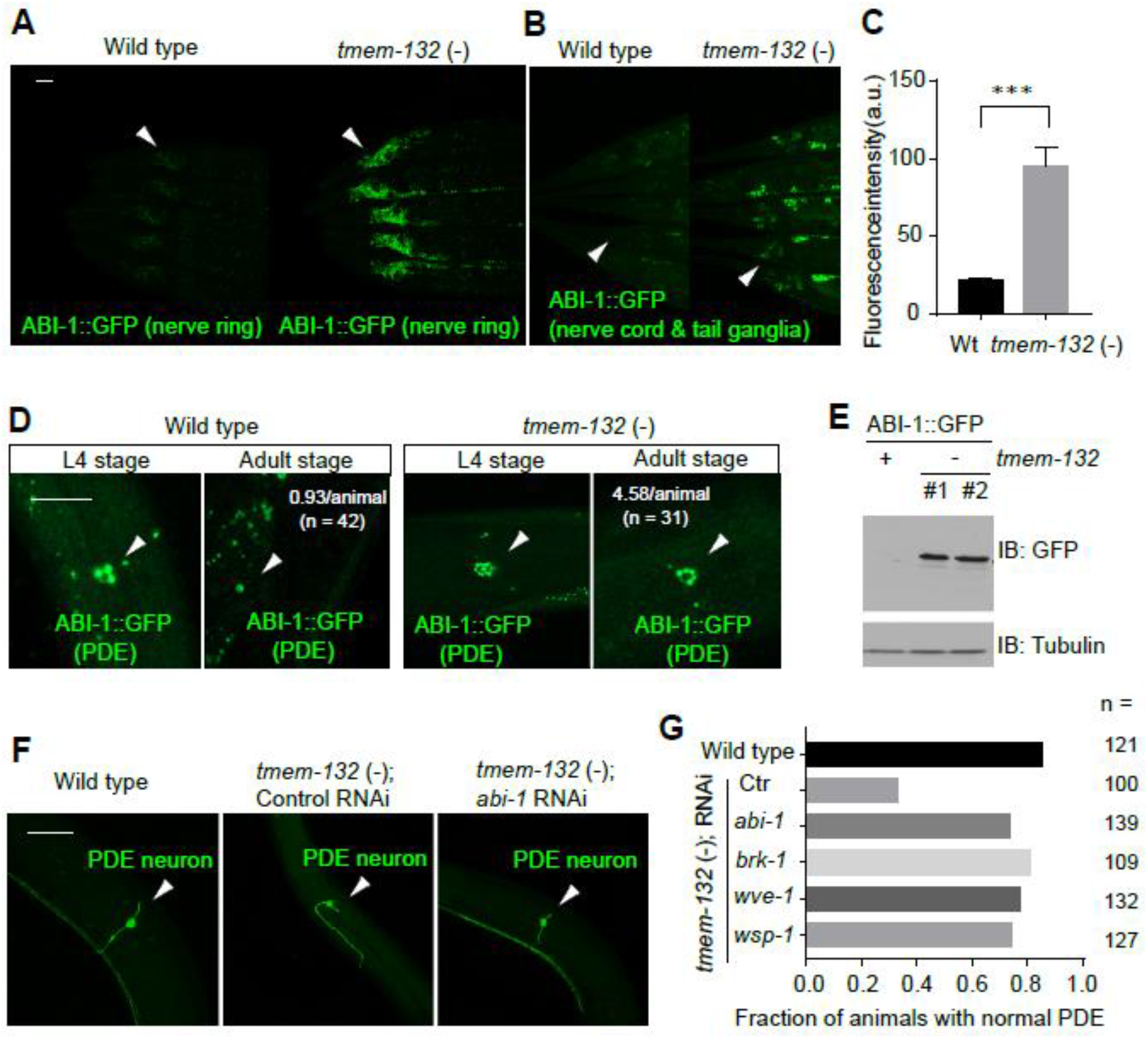
*C. elegans* TMEM-132 acts through WRC to regulate neuronal morphology. **(A)** Exemplar confocal fluorescent images showing increased abundance of neuronal ABI-1∷GFP in *tmem-132* deficient *C. elegans*. The *rab-3* promoter-driven expression of *abi-1*∷GFP was prominent in both nerve ring (arrow head in A) and tail ganglia (arrow head in B) areas of *tmem-132* null animals but not wild type. **(C)** Quantification of fluorescence intensity of ABI-1∷GFP in *tmem-132* null and wild-type animals. **(D)** Exemplar confocal fluorescent images showing increased abundance of ABI-1∷GFP in the PDE neurons of *tmem-132* null, compared with wild type animals. Up-regulation of ABI-1∷GFP in PDE was particularly prominent in young adult stage animals, compared with larval L4 stage animals. The average number of ABI-1∷GFP puncta were noted for adult stages. **(E)** Exemplar Western blot showing increased abundance of ABI-1∷GFP in total lysate of *tmem-132* null animals, compared with wild type. Two independent deletions (#1, *dma317*; #2, *dma318*) produced similar effects. **(F)** Exemplar confocal fluorescence images showing abnormal PDE morphology in *tmem-132* nulls, and rescued PDE morphology in *tmem-132* nulls with treatment of *abi-1* RNAi. **(G)** Quantification of fraction of animals with normal PDE morphology under indicated genetic conditions. *abi-1*, *brk-1*, *wsp-1* and *wve-1* encode components of WAVE or WAVE-like complex and their reduction-of-function by RNAi partially rescued abnormal PDE morphology of *tmem-132* null animals. Scale bars: 50 μm. *** indicates P < 0.001 (n = 5, repeated in at least three independent experiments).

### Ectopic TMEM132D expression decreases the abundance of select WRC components in mammalian cells

We next examined functional consequences of ectopic TMEM132D expression in mammalian cells. While WRC is present in most metazoan cell types, expression of endogenous *TMEM132D* appears to be limited to the nervous system, based on RNA profiling of various mammalian tissues (Figure S4). Similarly, we found that expression of *C. elegans tmem-132* localizes mostly, if not exclusively, in neurons (Figure 1B-D). We thus established a heterologous HEK293 cell line that stably expresses exogenous V5 epitope-tagged TMEM132D to assess how ectopic expression of TMEM132D affects abundance of WRC components, actin cytoskeletal dynamics and cell motility. Quantified fluorescence signal and Western blot analyses revealed that TMEM132D-expressing cell lines markedly decreased the abundance of ABI1∷GFP and WAVE2∷GFP while not apparently affecting that of NAP1∷GFP or SRA1∷GFP, after being transfected as GFP fusion constructs in control and TMEM132D cell lines (Figures 7A and 7B). NAP1 and SRA1 are essential for WRC stability and preventing other WRC components from degradation in the cell (Davidson et al., 2013; Eden et al., 2002; Kunda et al., 2003). Specific down-regulation of ABI1∷GFP and WAVE2∷GFP but not NAP1∷GFP or SRA1∷GFP suggests that TMEM132D likely acts to sequester NAP1 and SRA1 in a sub-complex from WRC, leading to disintegration and thus decreased activity of WRC.

**Figure 7.**
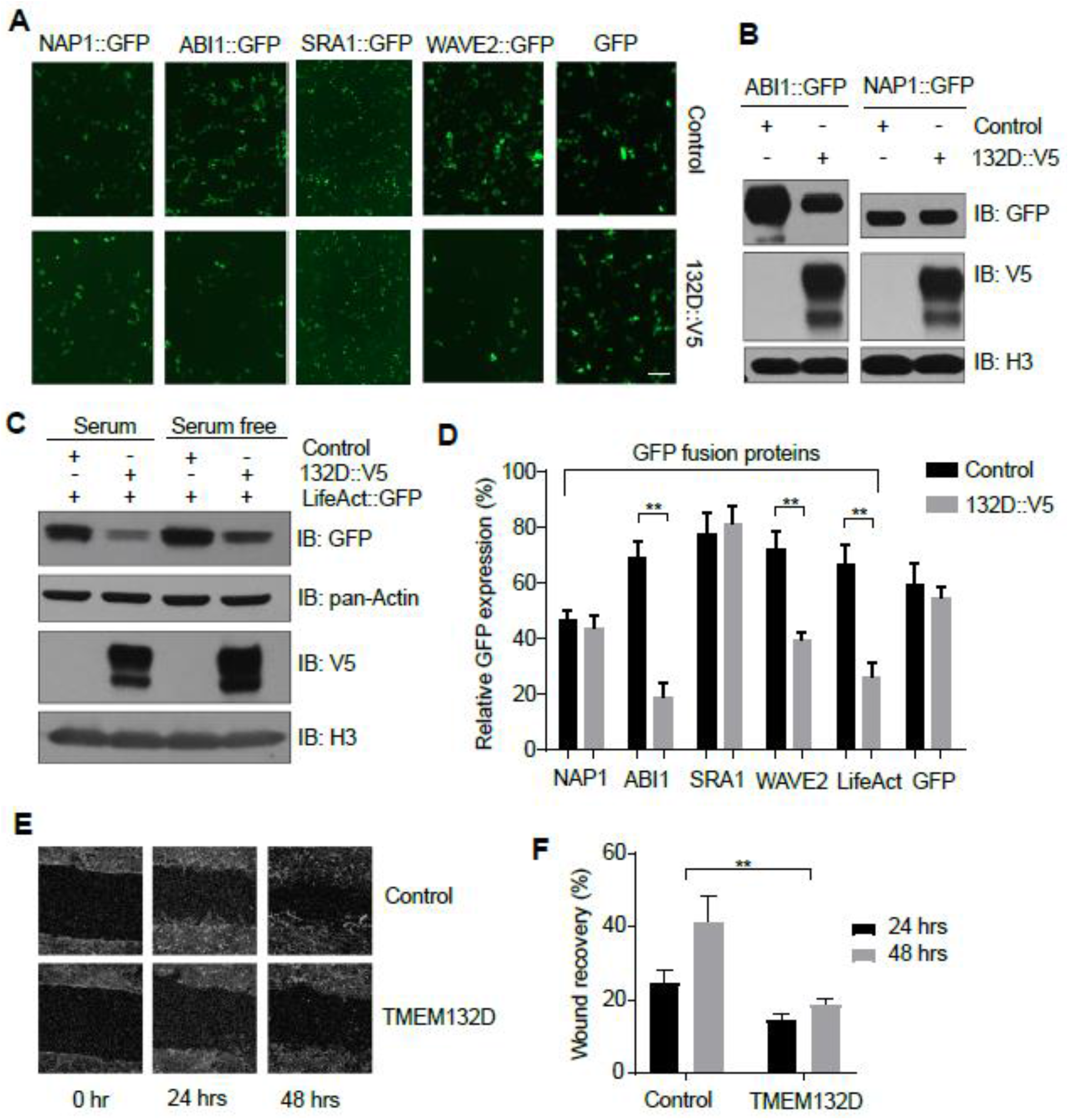
TMEM132D decreases abundance of selective WAVE components. **(A)** Exemplar epifluorescence images showing effects of TMEM132D overexpression on abundance of GFP-tagged WAVE components in HEK293 cells. Expression constructs encoding TMEM132D and individual components of WRCs or GFP only control were co-transfected for expression and imaging at 48 hrs post transfection, followed by quantification of percentage of GFP+ cells as shown in Figure 2D. **(B)** Exemplar Western blot showing decreased abundance of ABI1 but not NAP1 by co-expression with V5-tagged TMEM132D. Comparable abundance of V5 and H3 controls effects of TMEM132D expression levels and sample loading. **(C)** Exemplar Western blot showing decreased abundance of LifeAct∷GFP reporter by co-expression with V5-tagged TMEM132D, under serum-containing and serum-free media conditions. Comparable abundance of V5, pan-Actin and H3 controls effects of TMEM132D expression levels, monomeric Actin and sample loading respectively. **(D)** Quantification of percentage of GFP+ cells under indicated co-transfection conditions. **(E)** Exemplar micrographic images showing motile recovery of HEK293 cells stably expressing control or TMEM132D after line wounding. **(F)** Quantification of cell-free line width indicating wound recovery in HEK293 cells stably expressing control or TMEM132D 24 and 48 hrs after line wounding. Scale bars: 10 μm. ** indicates P < 0.01 (repeated in at least three independent experiments).

To test the prediction of the idea that TMEM132D inhibits WRC, we used the LifeAct reporter and a wound-recovery assay to examine effects of *TMEM132D* expression on actin nucleation and cell motility, respectively. LifeAct is a 17-amino-acid polypeptide that labels filamentous actin (F-actin) structures; its fusion with GFP allows visualization and quantification of actin nucleation in eukaryotic cells (Riedl et al., 2008a). We found that expression of epitope-tagged *TMEM132D* in HEK293T cells led to cell surface localization and markedly reduced the abundance of LifeAct∷GFP (Figures 7C, 7D and S5). This was the case even under the condition of serum starvation, which can increase LifeAct∷GFP abundance compared with the serum-containing condition (Figure 7C). Since a constitutive CMV promoter drove the expression of LifeAct∷GFP, altered abundance of LifeAct∷GFP likely reflects endogenous F-actin levels as unbound LifeAct∷GFP is unstable and likely degraded (Kumari et al., 2020; Riedl et al., 2008b). TMEM132D did not affect overall actin abundance based on Western blot analysis using a pan-actin antibody, indicating specific inhibitory effects of *TMEM132D* expression on F-actin but not actin monomers. In addition to overall abundance, quantitative cell population-level analysis revealed that TMEM132D also reduced the percentage of cells with strong LifeAct∷GFP fluorescence while not affecting the percentage of cells with control GFP fluorescence (Figure 7D). Furthermore, a wound-recovery assay showed that ectopic *TMEM132D*-expressing cells exhibited strongly reduced motility during the 24 and 48 hrs recovery phases after scratching-induced wounding in cultured cells (Figures 7E and 7F). Together, these results indicate that ectopic *TMEM132D* expression decreases actin nucleation and cell motility, supporting TMEM132D as a NAP1-binding and WRC-inhibiting protein.

## DISCUSSION

Bioinformatic analysis predicted a non-canonical cellular adhesion function for TMEM132 family proteins, connecting extracellular matrix with intracellular actin cytoskeleton (Sanchez-Pulido and Ponting, 2018). We provide experimental evidence to support this prediction and demonstrate that two members of the TMEM132 protein family from humans and *C. elegans* regulate cell motility and neuronal morphology, respectively, via inhibition of WRC and actin nucleation. Together, our data support a model in which TMEM132 family proteins via their C-termini bind to and sequester Nap/Sra away from WRC, leading to disintegration and decreased abundance/activity of WRC components such as Abi/Wave, and eventually reduced level of actin nucleation in the cell (Figure 8). Consequently, the high abundance of TMEM132 at local cell surface compartments in wild-type cells likely endows cells, including neurons, with restricted cell motility or morphogenesis, while deficiency of TMEM132 proteins may lead to development of inappropriate cell motility or ectopic morphogenesis. The extracellular part of TMEM132 family proteins contain three tandem immunoglobulin domains and a cohesin domain homolog with roles implicated in cellular adhesion (Sanchez-Pulido and Ponting, 2018). How TMEM132 proteins are regulated under physiological and pathological conditions, potentially through modulation by unidentified extracellular ligands, warrants further investigation.

**Figure 8.**
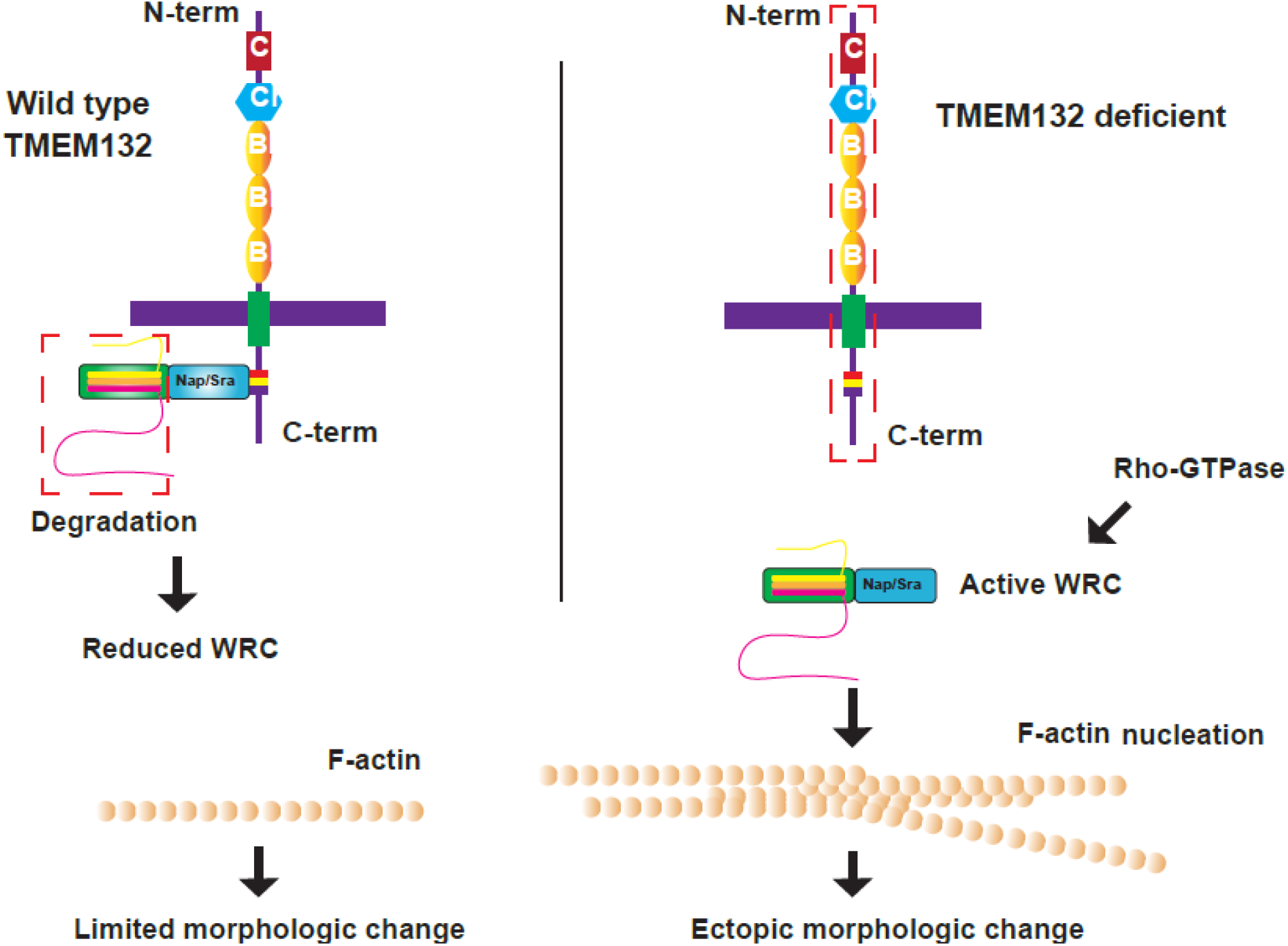
Model. In wild-type cells, TMEM132 family proteins function as local F-actin regulators by sequestrating Nap/Sra homologs and thus limiting the abundance of WRC because of disintegration/degradation of other WRC components. Consequently, there is limited cell morphological changes at cell surface compartments where TMEM132 proteins are enriched and WRC/F-actin nucleating activity is restricted. In TMEM132 deficient cells, Nap/Sra proteins are not sequestered and thus WRC is intact, permitting WRC/F-actin-nucleating activity that is normally regulated by Rho-GTPases and other factors in response to membrane signaling.

While major cell morphogenetic events occur during development, both human *TMEM132D* and *C. elegans tmem-132* are also highly expressed in mature neurons of the adult nervous system (Figures 1B-D and S4). Expression of the mouse homolog of *TMEM132D* is particularly high in the anterior cingulate cortex and claustral neurons, characteristic of long-range connectivity and cellular morphologic complexity (Erhardt et al., 2011; Saunders et al., 2018). Similarly, PDE neurons in *C. elegans* at the lateral side of the posterior body send long-range and bifurcated axons to the anterior and posterior nerve ganglia. Neurons form synaptic connections in circuits, in which neuronal activity dynamics can induce local F-actin-dependent changes of neuronal morphology and connectivity (Bertling and Hotulainen, 2017; Dillon and Goda, 2005; Luo, 2002). Such local changes are highly regulated while most surface compartments of mature neurons remain morphologically stable, mechanically resilient and maintained by large repertoires of cell adhesion molecules (Diz-Muñoz et al., 2018; Shapiro et al., 2007; Zipursky and Sanes, 2010). Correspondingly, enrichment of F-actin and actin-nucleating activity are also highly localized and differentially regulated along specific areas of cell processes and neuronal extensions, including dendritic spines, axonal synaptic termini, sensory cilia and microvilli ends (Balasanyan et al., 2017; Bertling and Hotulainen, 2017; Dillon and Goda, 2005; Drummond et al., 2018; Luo, 2002; Willig et al., 2014). We propose that TMEM132 family proteins act to restrict excessive WRC and actin nucleation activities, spatiotemporally necessary for cellular/neuronal morphological plasticity and maintenance. Dysfunction or dysregulation of TMEM132D may lead to abnormal neuronal structure and dynamics, contributing to heightened risks for depression, anxiety and panic disorders.

## Materials and Methods

### *C. elegans* strains

*C. elegans* strains were maintained with standard procedures unless otherwise specified. The N2 Bristol strain was used as the reference wild type. The genetic and transgenic alleles described in this study include Chr. III: *tmem-132*(*dma313*), *tmem-132*(*dma317*), *tmem-132*(*dma318*), *tmem-132*(*dma319*), *tmem-132*(*dma348*); *dmaEx471* [*tmem-132*p∷*tmem-132*fl∷GFP]; *lqIs2* [*osm-6*∷GFP]; *dmaEx452* [*osm-6*p∷*abi-1*∷GFP; *unc-54*p∷mCherry]; *dmaIs65* [*rab-3*p∷*abi-1*∷GFP; *unc-54*p∷mCherry]; *dmaIs86* [*osm-6*p∷*abi-1*∷GFP; *unc-54*p∷mCherry]; *dmaIs91* [*osm-6*p∷*tmem-132*∷GFP]; *wyIs592* [*ser-2*p3∷myr-GFP; *odr-1*p∷mCherry]; *otIs181* [*dat-1*∷mCherry + *ttx-3*∷mCherry]; *kyIs140* [*str-2*∷GFP + *lin-15*(+)].

### Yeast two hybrid assay and screen

The cDNA coding sequence of the C-terminal domain of human TMEM132D was cloned into the pGBKT7 vector and screened with a normalized universal human cDNA library (Clontech, 630481) in pGADT7 Vector, following instructions in the Matchmaker^®^ Gold Yeast Two-Hybrid System (Clontech, 630489). Verification of positive colonies was achieved by co-transformation of extracted bait and prey plasmids following the instruction of YeastMaker™ Yeast Transformation System 2 (Clontech, 630439) and by bait/prey plasmids with re-cloned cDNA.

### Co-immunoprecipitation and Western blot

HEK293T cells transfected with mammalian expression plasmids were pelleted by centrifugation, washed once with ice-cold PBS and lysed on ice for 30 min in lysis buffer (50 mM Tris HCl pH 8, 150 mM NaCl, 0.75% NP-40, 0.5% sodium deoxycholate) or Cell Lysis Buffer (Cell Signaling Technology, 9803S) supplemented with protease inhibitor cocktail (Sigma, 11836153001) and phosphatase inhibitor cocktail (Bimake, B15001). Following centrifugation at 12,000 rpm at 4°C for 15 min, supernatants were recovered. 10% volume of whole cell lysates were collected as input. Lysates were incubated with control (Chromotek) or rabbit IgG beads (Fisher Scientific, 88802) for preclear at 4°C for 45 min. Supernatants were recovered and incubated with mCherry-Trap, GFP-Trap (Chromotek) or V5 magnetic beads (MBL International, M167-11) at 4°C for 2 hrs. The beads were washed five times by lysis buffer and boiled with SDS sample buffer (Bio-rad, 1610747), then separated on 4-15% SDS-PAGE gel (Bio-Rad, 4561086) together with input. The proteins were transferred to a nitrocellulose membrane (Bio-Rad, 1620167) and detected using the GFP (Santa Cruz Biotechnology, sc-9996) or V5 (EMD Millipore, AB3792) antibody.

### Confocal and epifluorescence microscopic imaging

SPE confocal (Leica) and digital automated epifluorescence microscopes (EVOS, Life Technologies) were used to capture fluorescence images. Animals were randomly picked at the same stage and treated with 1 mM Levamisole sodium Azide in M9 solution (31742-250MG, Sigma-Aldrich), aligned on an 4% agar pad on a slide for imaging. Identical setting and conditions were used to compare experimental groups with control. For quantification of GFP fluorescence animals were outlined and quantified by measuring gray values using the ImageJ software. The data were plotted and analyzed by using GraphPad Prism7.

### Mammalian cell culture and wound recovery assay

U2OS and HEK293T cells were cultured in DMEM (Thermo Fisher Scientific, MT-10-013-CV), supplemented with 10% fetal bovine serum (FBS, Gemini Bio-Products, 900-208) and 1% penicillin/streptomycin in a humidified 5% CO2 incubator at 37°C. Stably expressing TMEM132D∷V5 or the control U2OS cells were constructed for wound recovery assay according to the reported protocol (Liang et al., 2007). Cells were plated onto the 6-well plate and grew for 16 hrs to create a confluent monolayer, then cells were washed once by DMEM and cultured in scratch medium (DMEM supplemented with 0.5% FBS and 1% penicillin/streptomycin) for 24 hrs. The cell monolayer was scraped in a straight line to create a “scratch” with a P200 pipet tip. The debris was removed by washing the cells three times with DMEM medium (0 hr). The cells were then cultured for 24 hrs in scratch medium and imaged at 0 hr, 24 hrs and 48 hrs. Human excitatory neurons were derived from inducible neurogenin-2(Ngn2) iPSC (i3N iPSCs) as described previously(Wang et al., 2017). Briefly, i3N iPSCs were pre-differentiated in KnockOut DMEM/F12 complemented with 2 mg/ml doxycycline, 1 mg/ml mouse Laminin, 10 ng/ml BDNF, 10 ng/ml NT3, 1x N-2 and 1x NEAA for 3 days. Media was changed daily with 10 mM Rock inhibitor added only for the first day. After that, the pre-differentiated precursor cells were disassociated with accutase and re-plated into poly-L-lysine coated plates in maturation media which is composed of DMEM/F12: Neurobasal-A/1:1, 2 mg/ml doxycycline, 1 mg/ml mouse Laminin, 10 ng/ml BDNF, 10 ng/ml NT3, 0.5x N-2, 0.5x B-27, 0.5x GlutaMax and 1x NEAA. Half of the media was replaced every week thereafter without doxycycline supplemented. Human excitatory neurons were infected with lenti-virus (MOI=1) for 24 hours on day 2 of pre-differentiation step. The precursor cells were then re-plated onto coverslips for differentiation into mature neurons and sample replicates were fixed every 3 days for immunohistochemistry.

### Statistical analysis

Data are presented as means ± S.D. with p values calculated by one-way or two-way ANOVA. Data with non-normal distribution, including gene expression and phenotypic penetrance results, were assessed by nonparametric Mann-Whitney and Fisherxs’s exact test, respectively.

## Acknowledgements

We thank the *Caenorhabditis* Genetics Center and the Bargmann, Hobert, Sengupta and Shen laboratories for various *C. elegans* strains, Dr. Orion Weiner’s laboratory at UCSF for WRC fluorescent reporter constructs, and Dr. Li Gan’s laboratory at the Gladstone Institute and UCSF for iPSC lines. The work was supported by NIH grants R01GM117461, Pew Scholar Award, Packard Fellowship in Science and Engineering (D.K.M), a fellowship from National Natural Science Foundation of China (X.W.) and a China Postdoctoral Foundation fellowship (W.J.).

## Author contributions

D.M., Y.S., Y.D. served as scientific advisors. X.W., W.J., S.L., B.W., X.Y., C.W. collected, analyzed and presented data. X.W., D.M., S.L., B.W. participated in writing and technical editing of the manuscript.

## Supplemental Figures and Table

**Supplemental Figure S1.**
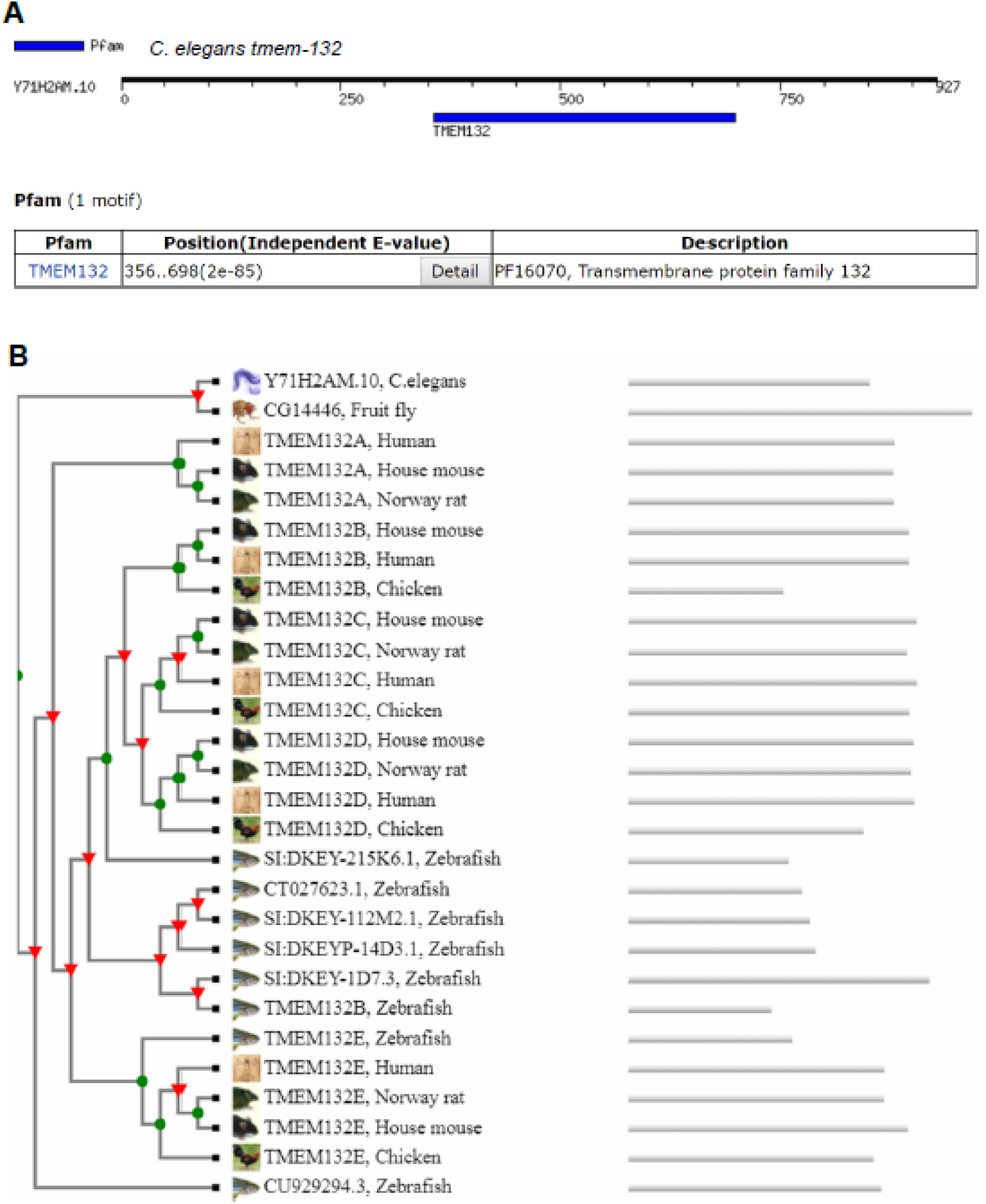
*C. elegans* TMEM-132 is a member of the evolutionarily conserved TMEM132 protein family. **(A)** Conserved Pfam domain in *C. elegans* TMEM-132 identified from MOTIF Search (https://www.genome.jp/tools/motif/MOTIF2.html). **(B)** Phylogenetic profile (generated by Wormbase.org) of the TMEM132 protein family with homologs from major metazoan species. Only *C. elegans* and fruit fly have one ortholog each.

**Supplemental Figure S2.**
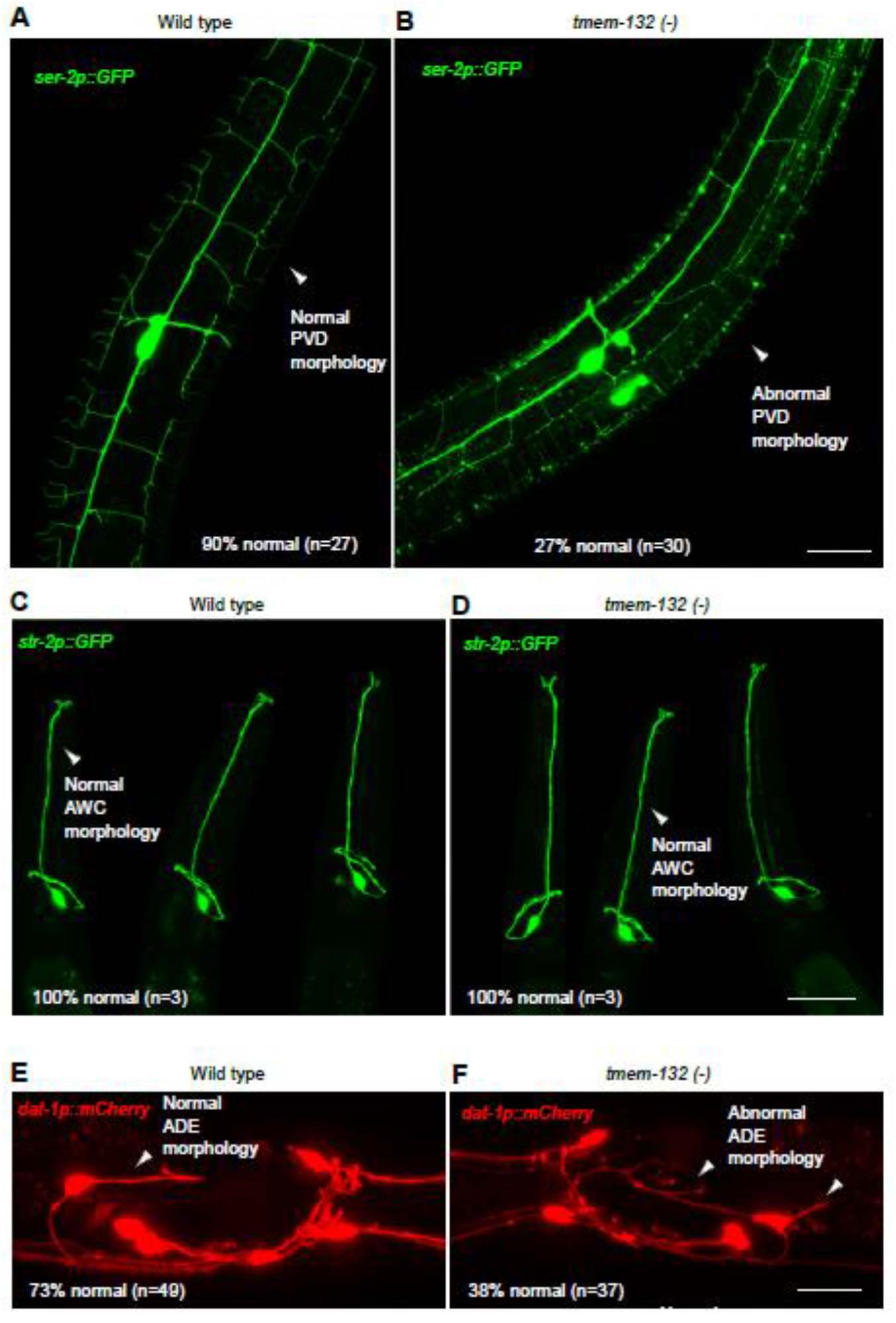
*C. elegans* TMEM-132 maintains morphologically complex PVD, ADE but not AWC neurons. **(A)** Exemplar confocal fluorescence image showing normal PVD neuronal morphology in wild type animals. **(B)** Exemplar confocal fluorescence image showing abnormal PVD neuronal morphology in *tmem-132* deletion mutants. **(C)** Exemplar confocal fluorescence image showing normal AWC neuronal morphology in wild type animals. **(D)** Exemplar confocal fluorescence image showing normal AWC neuronal morphology in *tmem-132* deletion mutants. **(E)** Exemplar confocal fluorescence image showing normal ADE neuronal morphology in wild type animals. **(F)** Exemplar confocal fluorescence image showing abnormal ADE neuronal morphology in *tmem-132* deletion mutants. Phenotypic penetrance (%) is noted.

**Supplemental Figure S3.**
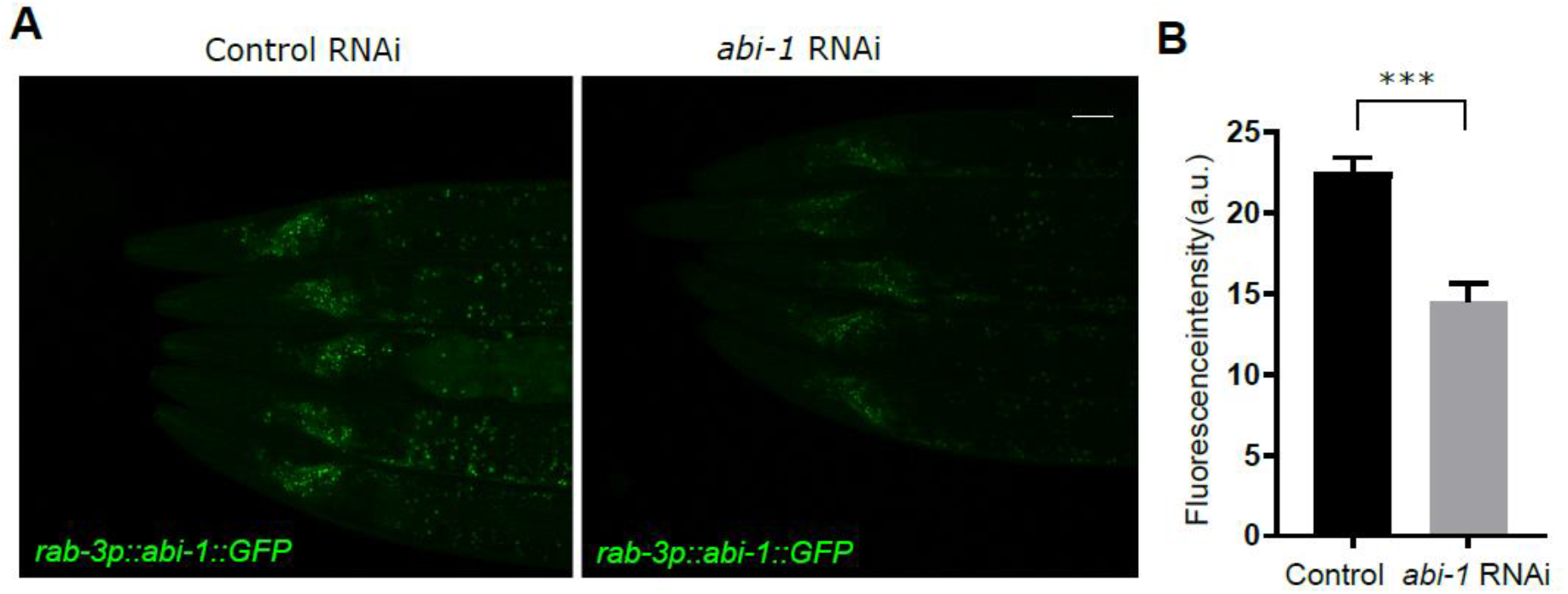
Efficacy of RNAi in neurons by feeding from bacteria. **(A)** Exemplar confocal fluorescence image showing *rab-3* promoter-driven expression of ABI-1∷GFP in control and animals fed with *E. Coli* expressing RNAi against *abi-1*. Scale bar: 50 μm. **(B)** Quantification of ABI-1∷GFP fluorescence intensity in control and animals fed with *E. Coli* expressing RNAi against *abi-1*. *** indicates P < 0.001 (n = 5, repeated in at least 3 independent experiments).

**Supplemental Figure S4.**
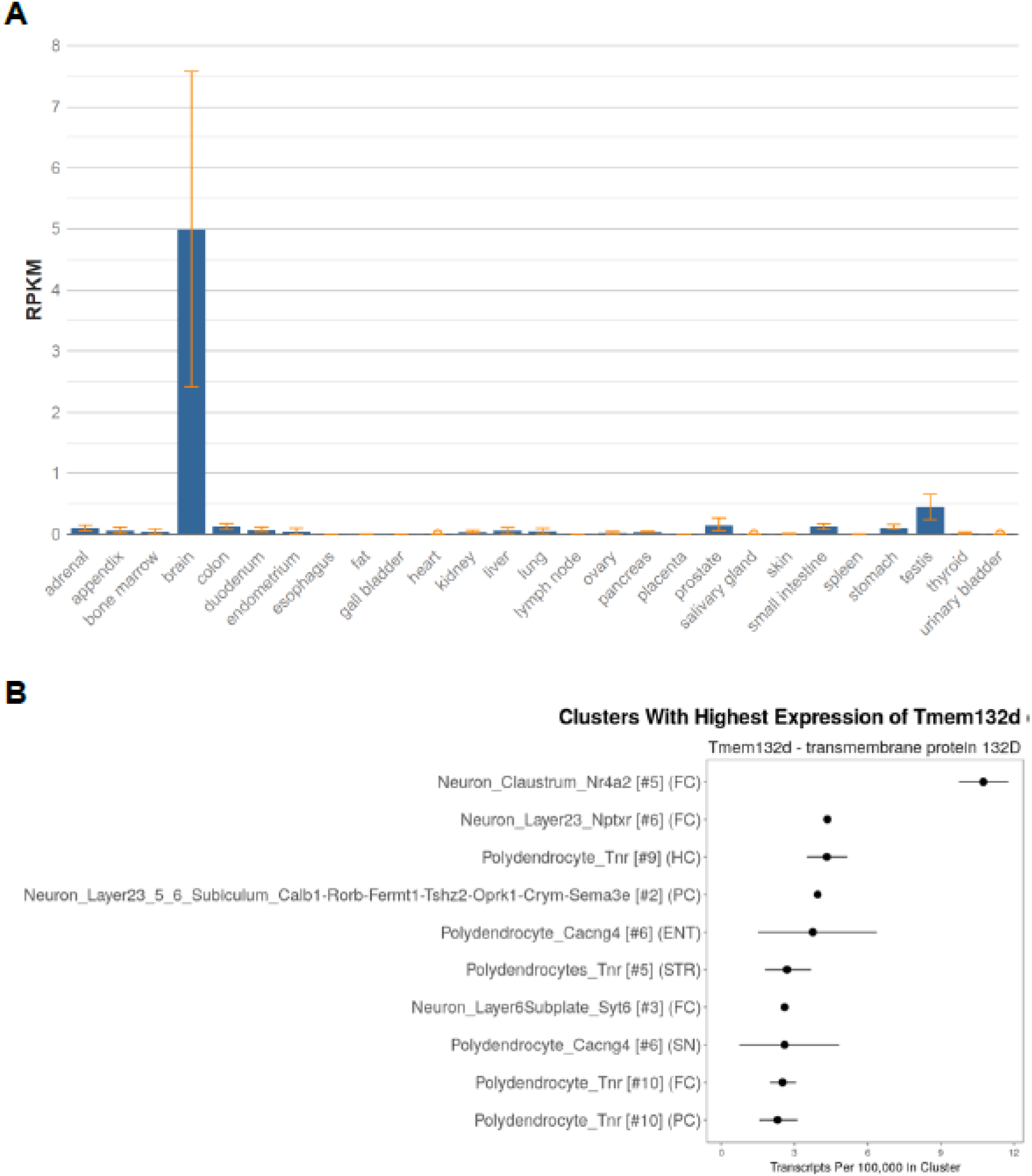
Enrichment of mammalian *TMEM132D* expression in the brain and claustral neurons. **(A)** NCBI graph (https://www.ncbi.nlm.nih.gov/gene/121256/?report=expression) quantification of human TMEM132D transcript abundance across 27 different tissues showing enrichment in brain. **(B)** DropViz (http://dropviz.org/) graph showing enrichment of *tmem132d* expression in mouse claustral neurons based on transcript abundance.

**Supplemental Figure S5.**
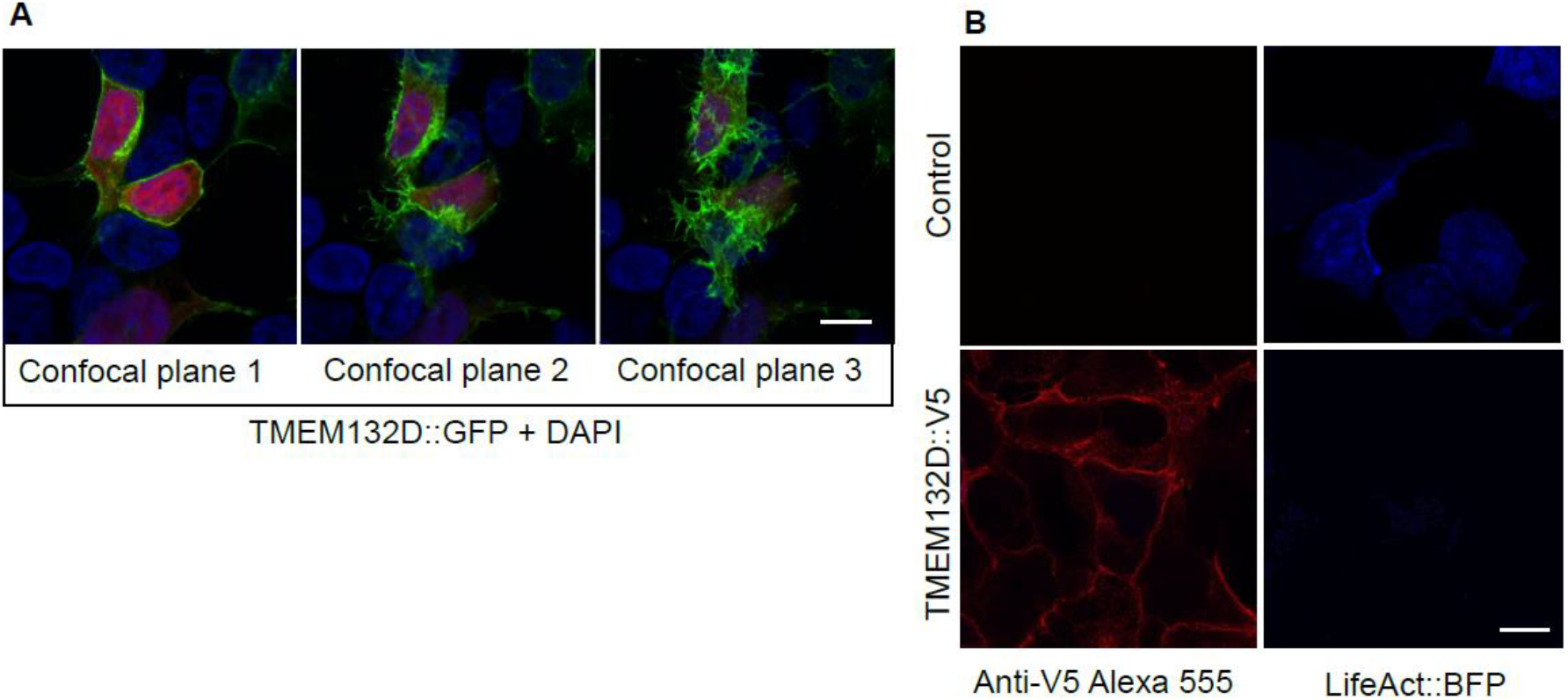
TMEM132D regulates F-actin abundance. **(A)** Exemplar confocal fluorescence images (from three different confocal planes) showing membrane localization pattern of TMEM132D∷GFP in HEK293 cells. **(B)** Exemplar confocal fluorescence images showing immunostaining of V5-tagged TMEM132D and BFP-tagged LifeAct in HEK293 cells. V5 positive cells strongly decreased abundance of LifeAct∷BFP. Scale bar: 10 μm.

**Supplementary Table 1.**
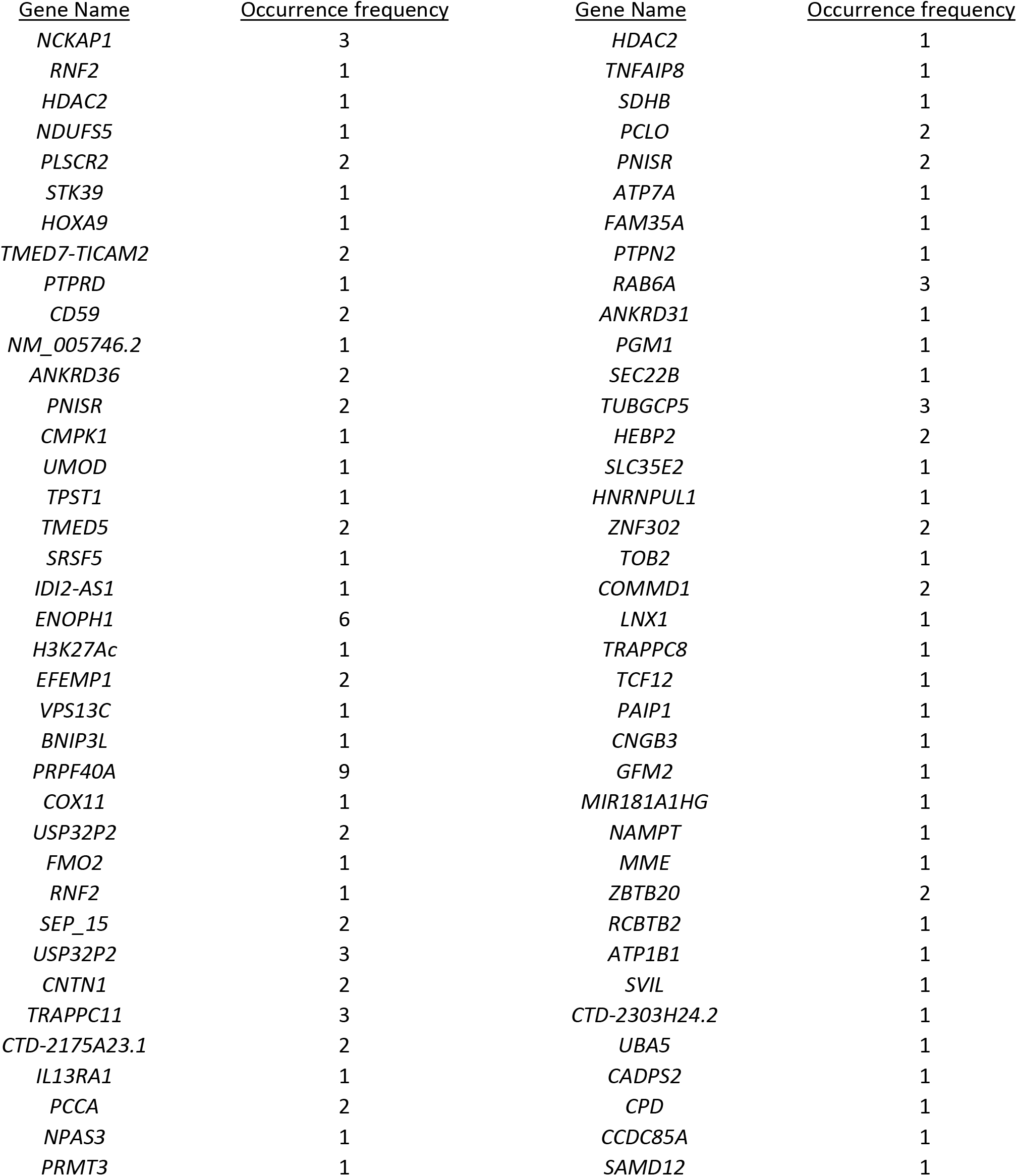
List of TMEM132D interactor-encoding genes identified from yeast-two-hybrid screens.

## Notes

### Competing Interest Statement

The authors have declared no competing interest.

